# WhatIsMyGene: Back to the Basics of Gene Enrichment

**DOI:** 10.1101/2023.10.31.564902

**Authors:** Kenneth Hodge, Thammakorn Saethang

**Affiliations:** Unaffiliated; Department of Computer Science, Faculty of Science, Kasetsart University, Bangkok, 10900, Thailand

## Abstract

Since its inception over 25 years ago, gene enrichment has been largely associated with curated gene lists (e.g. GO) that are constructed to represent various biological concepts: the cell cycle, cancer drivers, protein-protein interactions, etc. Researchers expect that a comparison of their own lab-generated lists with curated lists should produce insight. Despite the abundance of such curated lists, we here show that they rarely outperform comparisons against existing individual lab-generated datasets when measured using standard statistical tests of study/study overlap. This demonstration is enabled by the WhatIsMyGene database, which we believe to be the single largest compendium of transcriptomic and micro-RNA perturbation data. The database also houses voluminous proteomic, cell type clustering, lncRNA, and epitranscriptomic data. In the case of enrichment tools that do incorporate specific lab studies in underlying databases, WIMG generally outperforms in the simple task of reflecting back to the user known aspects of the input set (cell type, the type of perturbation, species, etc.), enhancing confidence that unknown aspects of the input may also be revealed in the output. To optimize the ranking of gene enrichment results, WIMG imputes backgrounds to curated gene lists, an approach we investigate with Shannon information analysis. Thus, gene lists that are replete with abundant entities do not inordinately percolate to the highest-ranking positions in output. We delineate a number of other features that should make WIMG indispensable in answering essential questions such as “What processes are embodied in my gene list?” and “What does my gene do?”

## Main

Modern transcriptomic protocols typically result in the identification of 20,000 RNA species. Proteomic datasets may now contain over 8,000 proteins. In most cases, researchers are interested in subsets of identified entities, e.g. the set of transcripts that are strongly up-regulated when interferon is applied to a particular cell line. Once such a subset has been selected, the researcher seeks to compare the subset with curated gene lists (CGLs) that presumably reflect important biological processes. In the case of the interferon experiment, it is likely that a CGL with a term such as “innate immunity” would best match the experimental subset (ES). In many cases, the researcher may have little expectation as to the result of such a comparison. A great deal of effort has been devoted to the construction of curated lists. At the time of writing, the GO consortium[1] offers 42,887 terms (lists), with KEGG[2], PANTHER[3], REACTOME[4], GSEA (via “Hallmark” gene sets[5]) and others, each offering thousands of terms.

CGLs represent biology’s best attempts to identify gene sets that recurrently appear over multiple experiments. However, problems may arise when various statistical tests are applied in order to isolate and rank the CGLs that best match an experimental result. These issues include the potential absence of important genes in CGLs[6], the over-representation of particular genes over many CGLs[6–9], derivation of CGLs from a small subset of relevant studies and cell types[6, 10], significant changes in the composition of CGLs over time (the “reproducibility crisis”[8]), absence of human oversight in generating CGLs[8], distortion due to the effects of multifunctional genes in CGLs[11], and the questionable assumption that biological entities are independently expressed[12, 13]. The blame for some of these deficiencies need not rest solely on the curating bodies, as experimentalists may choose to research predictable or “hot” genes in pursuit of job security and other mundane concerns[14–16]. To the above concerns we would add the possibility that the typical ES is not best described by the themes that are easily subsumed by CGLs[17], but rather, in terms of modules of genes that do not share obvious functions. A final important concern is the simple fact that CGLs lack backgrounds. That is, the complete set of genes from which a CGL subset was drawn, a critical input for statistical analysis, is unknown. Typically, analysis tools operate under the assumption that a CGL’s background is simply the total number of known genes for an organism[18–21], or the total number of genes found in all CGLs[22, 23]. Gene enrichment tutorials may emphasize that the background of the experimental subset must be carefully considered, but seem unconcerned with the fact that CGLs themselves do not have backgrounds.

Numerous attempts have been made to resolve the above concerns by generating improved algorithms for gene enrichment analysis. We promote a solution that resides not primarily in algorithms, but in the composition of the enrichment database: a “study/study” approach matching one’s own experimental subset with subsets generated by other experimenters[10]. RNA-seq results, for example, do not ordinarily suffer from over-representation of popular genes, the purposeful ignoring of rare or uninteresting genes, fads in the field, or unusual biases in the appearance of multifunctional genes. The experimental subsets identified via RNA-seq experiments are not expected to “evolve” over the years. Furthermore, an RNA-seq experiment’s background can be empirically ascertained. Below, we further explore some difficulties presented by the usage of CGLs, and detail how WhatIsMyGene, via sheer database size and a variety of analysis tools and approaches, may greatly aid in the generation of biological insight.

## Results

### The absence of CGL backgrounds is consequential

It could be argued that the decision to assume either a large or small background for a CGL would not have significant repercussions on output. That is, when ranking the relevance of various CGLs against an experimental subset, rankings will not change, or will change insubstantially, depending on the selection of CGL backgrounds.

We first illustrate the dependence of output *P*-values on CGL backgrounds. We use Fisher’s exact test for this exercise, but similar outcomes could be produced by any of the methods commonly used by ORA (over-representation analysis) tools. This test requires four inputs: 1) the number of genes in the ES, 2) in the CGL, 3) within the ES/CGL intersection, and 4) found within the intersection of ES and CGL backgrounds. Given 4), if the ES background is large relative to the CGL background, the fourth input cannot be larger than that of the CGL background. In Fig. 1A, four combinations of these four inputs are used to generate curves representing output *P*-values, with only the CGL background allowed to vary. For simplicity, the ES size and the CGL size are equal in all cases. In the image, case 2 illustrates a “typical” ES/CGL result, where significance is seen only when the actual intersection is considerably less than the expected intersection. Such a result would usually be excluded from a supplemental dataset of ranked CGLs. Case 1 is of greater interest. Here, if the CGL has a background greater than 8600, the ES/CGL intersection would be significant. However, a CGL background below 4600 would also be significant due to anti-correlation between ES and CGL. One may imagine a case where the ES represents genes down-regulated on a particular drug treatment, and the CGL involves genes upregulated in Alzheimer’s disease. If the CGL “ground-truth” background is 20,000, an Alzheimer’s treatment may have been identified. However, if the CGL ground-truth background is 4,000, the drug treatment may actually accelerate the progression of Alzheimer’s disease. Comparing case 3 and case 4, case 4 settings initially outperform case 3 settings due to the presence of one extra gene in the case 4 ES/CGL intersection. However, the case 4 curve levels off at the background setting of the experimental dataset (10,000). In all cases, the choice of CGL background strongly influences the measure of significance.

**Figure 1.**
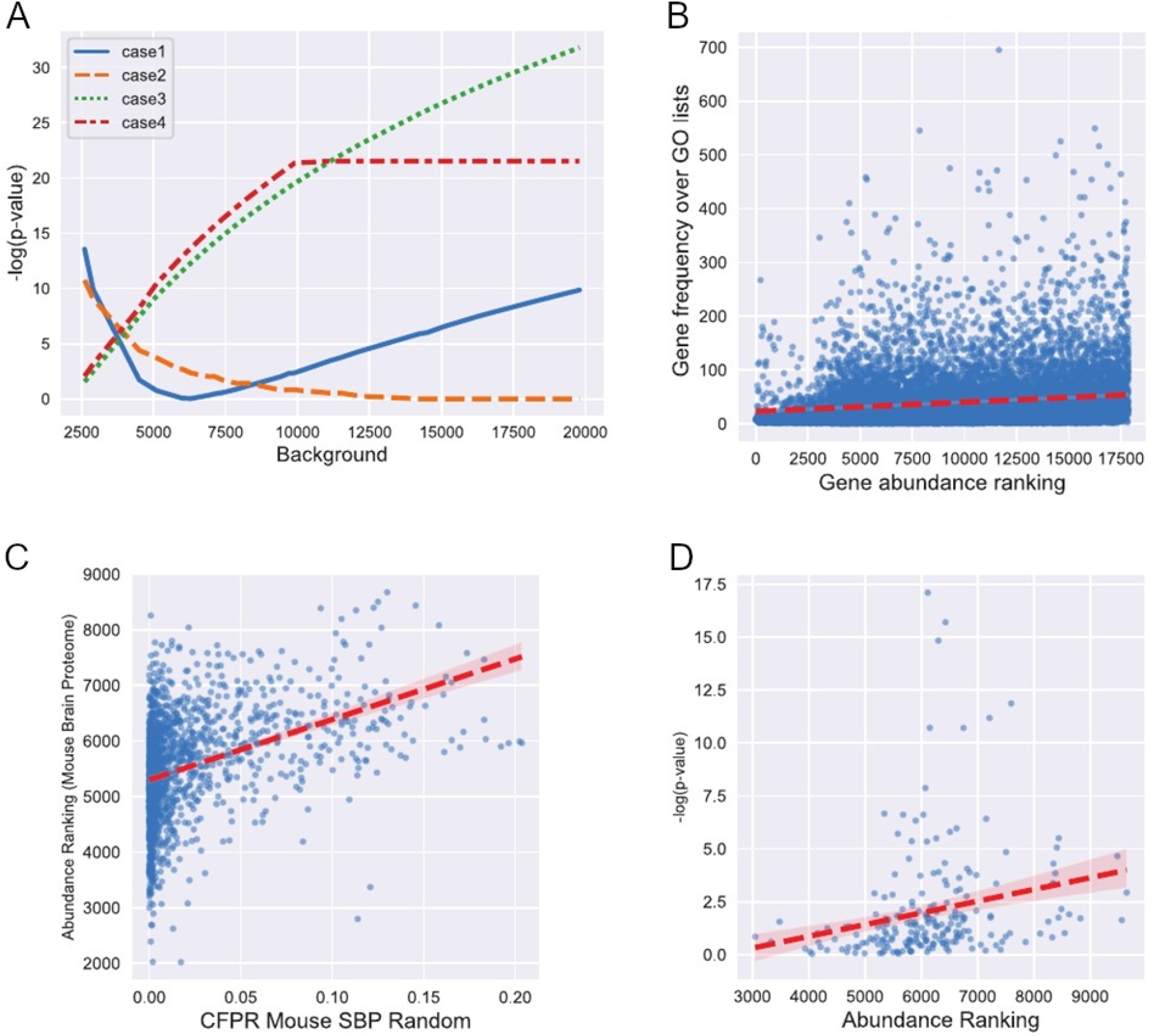
Assumptions regarding CGL background sizes may affect gene enrichment results. Fig 1A. Dependency of significance measurement on CGL background. Intersection, CGL size, ES size, and experimental background for Fisher’s exact test are as follows: Case 1 (40,500,500,20000), Case 2 (2,200,200,20000), Case 3 (20,200,200,20000), Case 4 (21,200,200,10000). The results assume that all genes comprising the CGL background are found in the experimental background. The possibility of testing over multiple CGLs is ignored in the significance calculations. Fig. 1B. Abundance vs frequency of GO-CGL genes. The frequency of gene appearance over 2604 GO-CGLs is plotted against abundance rankings derived from the whole human proteome; a high rank means high abundance (R^2^ = .155; *P* = 5.4*10^-73^). Fig. 1C. GO-CGL false positive rates correlate with transcriptomic abundance (R^2^ =.39, *P* = 1.6*10^-55^). Fig 1D. CGL enrichment tables may be biased toward abundance. 210 GO-CGLs are assigned *P*-values in a supplementary table. These values are plotted against abundance rankings for each GOCGL (R^2^ =.23, *P*= .0008). It is apparent that CGLs containing abundant genes tend to receive the most significant *P*-values.

One may ask how it is possible that the proper background for a CGL would be 4,000. We present several simple cases as illustrative of the possibility. First, consider two hypoxia-related studies, having backgrounds of 10,000 and 20,000 genes. Shall we assume that 10,000 genes are common to both sets? No, because the 10,000-member set may contain genes not found in the 20,000-member set. Perhaps the joint background is 9,000. We wish to include 10 more hypoxia studies as the basis for a CGL. It is apparent that incorporation of each additional experimental set lowers the background of the CGL. The general effect of this process is to enrich for abundant genes found in all datasets, and eliminate rare ones. We may now have a CGL with an effective background of 4,000. The problem is exacerbated by the inclusion of genes derived from studies with particularly small backgrounds, genes from “targeted microarrays”, or genes highlighted in studies that predate the big-data era.

Next, we generate a frequency table of genes found in 2,604 GO[1] CGLs. Using a whole-human list of proteins ranked according to abundance[24], frequencies of genes are plotted against their abundances (Table S1). Fig. 1B makes clear that abundant genes are indeed enriched in these CGLs; *P*= 5.4*10^-73^ (Pearson). Not surprisingly, the single most common gene over these CGLs is TNF, with 695 appearances. The gene is an outlier, however, given its moderate abundance ranking (the 6206^th^ most abundant gene in a list of 17861). Examples of genes that are both frequent and abundant are CTNNB1, RHOA, ANXA1, and HSPA1A.

Assessing GO-CGLs according to abundance rankings needn’t be limited to ES/GO-CGL combinations. Fulcher et al ranked GO-CGLs according to their tendencies to produce high “category false positive rates”[12] against mouse and human transcriptomic data. Using a list of mouse brain proteins ranked according to abundance[25], we first generate “abundance rankings” for 2604 GO-CGLs. The rankings reflect the mean abundance of genes in each CGL; CGLs with highly abundant genes receive high rankings. Fig. 1C shows a strong relationship between GO-CGL false-positive rates and our estimates of GO-CGL abundance (Table S2).

Finally, we invoke a supplemental table from an actual study in which the mouse brain proteome was measured after RAB3GAP1 disruption[26]. We plot the supplementary table *P*-values representing GO-CGL enrichment against our GO-CGL abundance rankings (Table S3). Fig 1D shows that GO-CGLs with high abundance rankings strongly correlate with the most significant *P*-values. An exhaustive survey of supplementary gene enrichment tables is beyond the scope of this work. Fulcher et al[12], in fact, performed this exercise for 31 brain transcriptomic studies, finding excessive representation of particular GO-CGLs over multiple phenotypes. The fact that multiple enrichment tools were used over these 31 studies suggests that bias, to some extent, lies outside the mere algorithms utilized by these tools. One should consider the possibility that numerous such tables are skewed in this way due to the absence of backgrounds for CGLs.

It might be instructive to examine the contents of a CGL with a particularly low abundance ranking. Numerous GO CGLs are devoted to ribosomal components; e.g. GO:0022625, “cytosolic large ribosomal subunit.” This CGL is indisputably an unabridged collection of large ribosomal components. Such genes are, of course, quite abundant over all cell types and organisms. Shall we, however, assign a background equivalent to the count of all genes in the genome to this CGL?

### Assigning backgrounds to gene sets

Whenever possible, WIMG records the “actual” background of genes involved in a study. If the study involves RNA-seq, for example, this is simply the count of all transcripts that pass quality control. Thus, the quality of any method that estimates the background of a set of genes can be assessed by comparing these estimates to actual backgrounds. Given the above construction of abundance rankings, it seems reasonable to hypothesize that CGL backgrounds could be estimated and then utilized as a more accurate input for Fisher’s exact test. This can be accomplished by simply doubling the averaged abundance rankings for all genes in a set (see Methods). Presumably, increased precision in these background estimates can be accomplished by creating multiple tissue and species-specific lists of proteins and transcripts ranked by abundance, to which ESs can then be compared.

We assigned backgrounds to 44,620 WIMG ESs based on cell type, species, and the gene composition of the individual ESs. Over these studies, we find high correlation between the actual backgrounds and the background implied by the gene composition (R^2^ =.126, P=6*10^-158^, Pearson). Fig. 2A plots the actual and implied background of a subset of 1% of these lists. The only requirement for entry into this test was that we had already generated abundance ranks for the specific cell types involved in the tests (e.g. an MCF7 study would require an MCF7-based list of genes ranked by abundance). Studies were entered as we encountered them in the GEO database or other sources. Another method of background estimation which we termed “reverse-Fisher” was also applied (Methods). Doubling of the abundance ranking predicted backgrounds with R^2^=.52, *P*=5*10^-21^, while the reverse-Fisher method gave R^2^=.54, *P*=4.7*10^-23^. We attribute the improvement in R values versus the initial large-scale test to proper matching of tissue types. Note that background prediction remained accurate even when the actual backgrounds were relatively small, as seen in proteomic datasets and occasional RNA-seq results.

**Figure 2.**
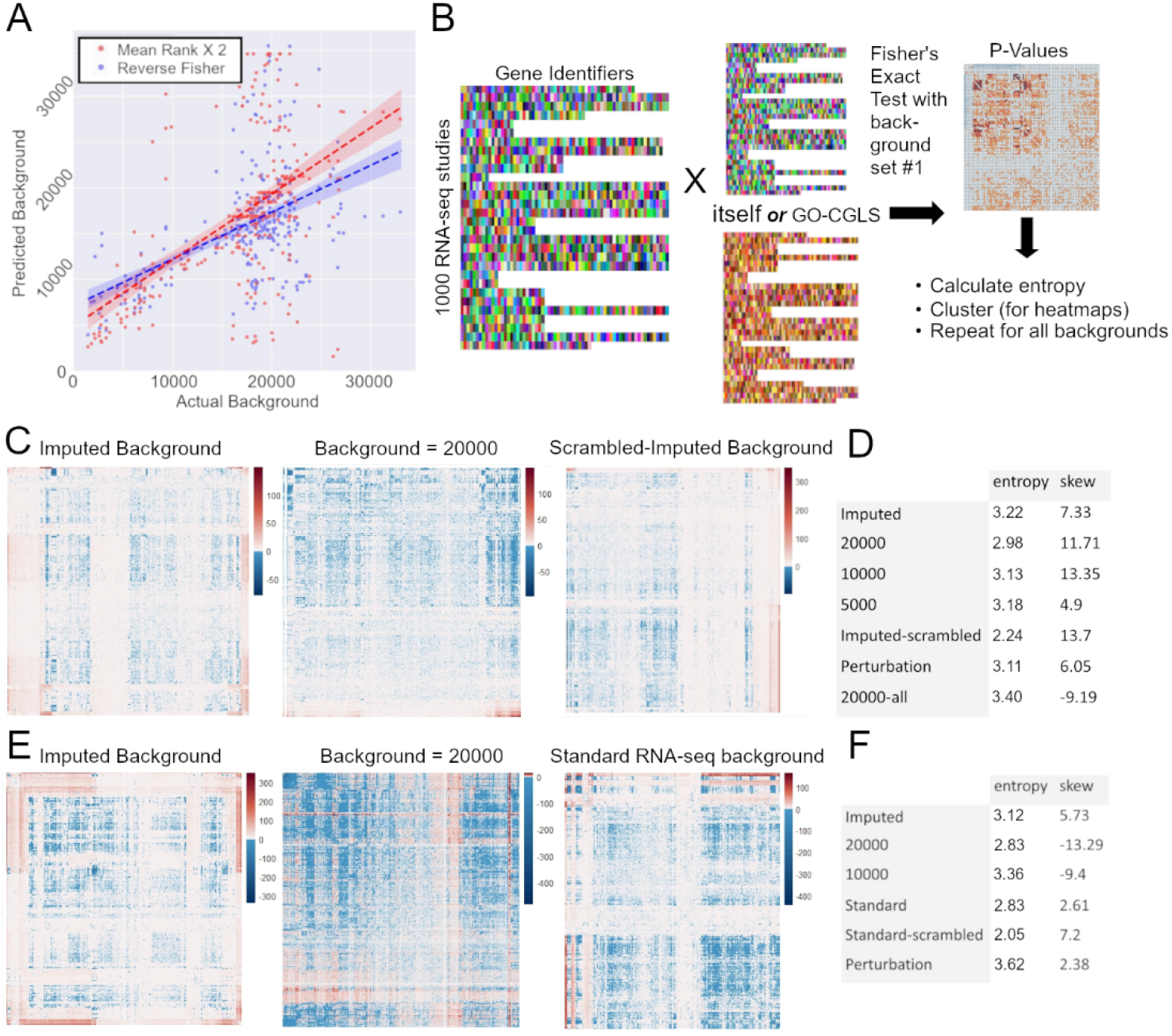
Validating Imputed Backgrounds. A. The actual backgrounds of 287 gene lists are compared against backgrounds estimated for these studies via a “reverse Fisher” approach and via doubling of mean abundance ranks. B. The scheme for comparison of various “backgrounding” approaches. *P*-values are derived for all combinations of RNA-seq/RNA-seq or RNA-seq/CGL lists. The *P*-values are then utilized for statistics and heatmaps. C. Clustering of RNA-seq/CGL *P*-values generated from three background approaches. In this scheme, negative values indicate set/set overlap; positive values correspond to absence. Note that, in addition to shifts in coloring and patterning, heatmap scales also change in response to background. D. Statistics for 7 CGL/ES background approaches. “20000-all” refers to a case in which a background of 20000 was applied not only to CGLs, but to RNA-seq lists as well; note the extreme negative skew in this case. E. Clustering of RNA-seq/RNA-seq *P*-values from three background approaches. F. Statistics for 6 ES/ES background approaches.

WIMG has proceeded to generate backgrounds for GO-CGLs and other gene lists (e.g. gene clusters generated from single-cell transcriptomic matrices). The assumption is that if background-adjustment offers reasonable background estimates for studies with known backgrounds, it should do the same for gene lists with unknown backgrounds. The median GO-CGL background was 10254 and the mean was 15300, with a low of 643 (GO:0072562, “blood microparticle”) and a high of 50148 (GO:0008237, “metallopeptidase activity”). Table S7 gives our estimated backgrounds for 2605 GO-CGLs. The underlying data is found in Table S9.

While the allure of extreme *P*-values is strong, we reason that an optimal gene-enrichment system should instead maximize the information content of its output. The resulting scores should typically exhibit neither a flat distribution of noise nor the excessive clustering indicative of systematic bias. We thus tested the effects of various background estimates on Shannon entropy. We randomly chose 1,000 human RNA-seq ESs and 1,000 GO CGLs, generating *P*-values for each of the 1,000,000 ES/CGL combinations under 7 background-estimate conditions (see Methods). Log(*P*-values) were signed; those generating higher than expectation intersections received negative values (as the log-value of a very significant *P*is strongly negative), while lower than expectation intersections received positive values. Shannon entropy was then calculated for each condition using standardized bins to ensure scale invariance (Fig. 2B-D). Here, imputed CGL backgrounds (average background 15,848) outperformed backgrounds uniformly set at 10,000 and the “industry standard” 20,000 with 6% and 18% more resolution, respectively. Though still underperforming imputation, merely setting all CGL backgrounds to 5,000 performed better than backgrounds set at 10,000 or 20,000, suggesting that CGLs are generally overloaded with abundant entities; we return to the subject of extreme low backgrounds in the next section. We also utilized “perturbation backgrounds”, which utilize underlying lists of genes ranked not by abundance, but by frequency of perturbation over the WIMG database. However, highest information content was obtained by setting both CGLs and ESs to a background of 20,000. Note that this biologically unrealistic approach also generated the only case of negative skew, with nearly 3% of all study/study intersections attaining significance after adjustment, compared to 0.5% for the second most information dense approach, that of imputation. Scrambling the backgrounds reduced entropy to the lowest observed levels, validating the metric’s ability to distinguish signal from noise.

In the case of ESs, whenever possible WIMG sets backgrounds according to the complete quality-controlled set of genes identified in a study (“standard” backgrounds). Nevertheless, it is likely that various algorithms that seek to rank differentially expressed genes (e.g. DESeq2) tend to emphasize abundant genes[27–29], potentially creating a discrepancy between the “effective” and “standard” backgrounds that not only biases results, but compresses the dynamic range of the output, effectively lowering the resolution of the statistical test. To determine if we could increase this discriminative power, we repeated our Shannon entropy analysis, this time examining the above set of 1,000 ESs crossed with the same 1,000 ESs (499,000 unique ES/ES intersections after excluding all intersections of an ES with itself) (Fig. 2E-F). Here, backgrounds reflecting perturbation frequency outperformed all other backgrounds in terms of information. The imputation approach outperformed a static background of 20,000, but was beaten by a static background of 10,000; both of these static backgrounds resulted in strongly negative skew, however. In fact, these two static backgrounds generated coefficients of kurtosis nearly triple those of other approaches, suggesting that these models fail to resolve gradations of biological significance, producing instead a binary distribution dominated by noise and punctuated by extreme, likely artifactual, outliers. Again, scrambling of backgrounds produced information content dramatically less than that of other approaches.

Complete entropy statistics are available in table S12.

### Benchmarking CGL backgrounds

If asked to design an experiment in which an ES strongly intersects with a randomly selected CGL, a biologist would likely find the task onerous; i.e. in most cases, it is not obvious what sorts of lab operations would be expected to generate an ES that matches well with a given CGL. For a minority of CGLs, however, it is fairly clear how a researcher would proceed. Such cases involve common perturbations. We selected a perturbation, the response to hypoxia, for which a CGL exists (GO:007082 Response to Oxygen Levels) and multiple relevant studies can be found in our database. When these studies are tested against 3719 CGLs, most of which are expected to be irrelevant to hypoxia, one would expect the hypoxia-relevant CGL to rank highly. We performed this test with 98 hypoxia studies, utilizing 6 fixed backgrounds as well as the set of backgrounds that WIMG imputes to these CGLs. This CGL indeed rose to rank #1 in 25 cases under at least one background setting. It is notable that in 59 of these studies, the assigned background outperformed all of the fixed backgrounds. Practically speaking, a biologist is unlikely to consider the possibility that hypoxia is relevant to an ES if no hypoxia-related CGL can be found within the set of highest-ranking CGLs. Excluding all studies in which a ranking of 20 or less was not found under any background setting left 49 studies. In 18 such cases, the WIMG background outperformed all other backgrounds; in 19 other cases, the WIMG background shared top ranking. Ignoring WIMG backgrounds, it is also notable that a background setting of 10000 produced 16 cases that obtained or shared highest rank; in no cases did a background of 25000 cause a study to obtain highest rank. Other trends, such as a paucity of cases that achieved highest ranking at a background of 15000 and a tendency for rankings to rise steeply at background settings less than that at which top ranking is achieved, are seen in Fig. 3B.

**Figure 3.**
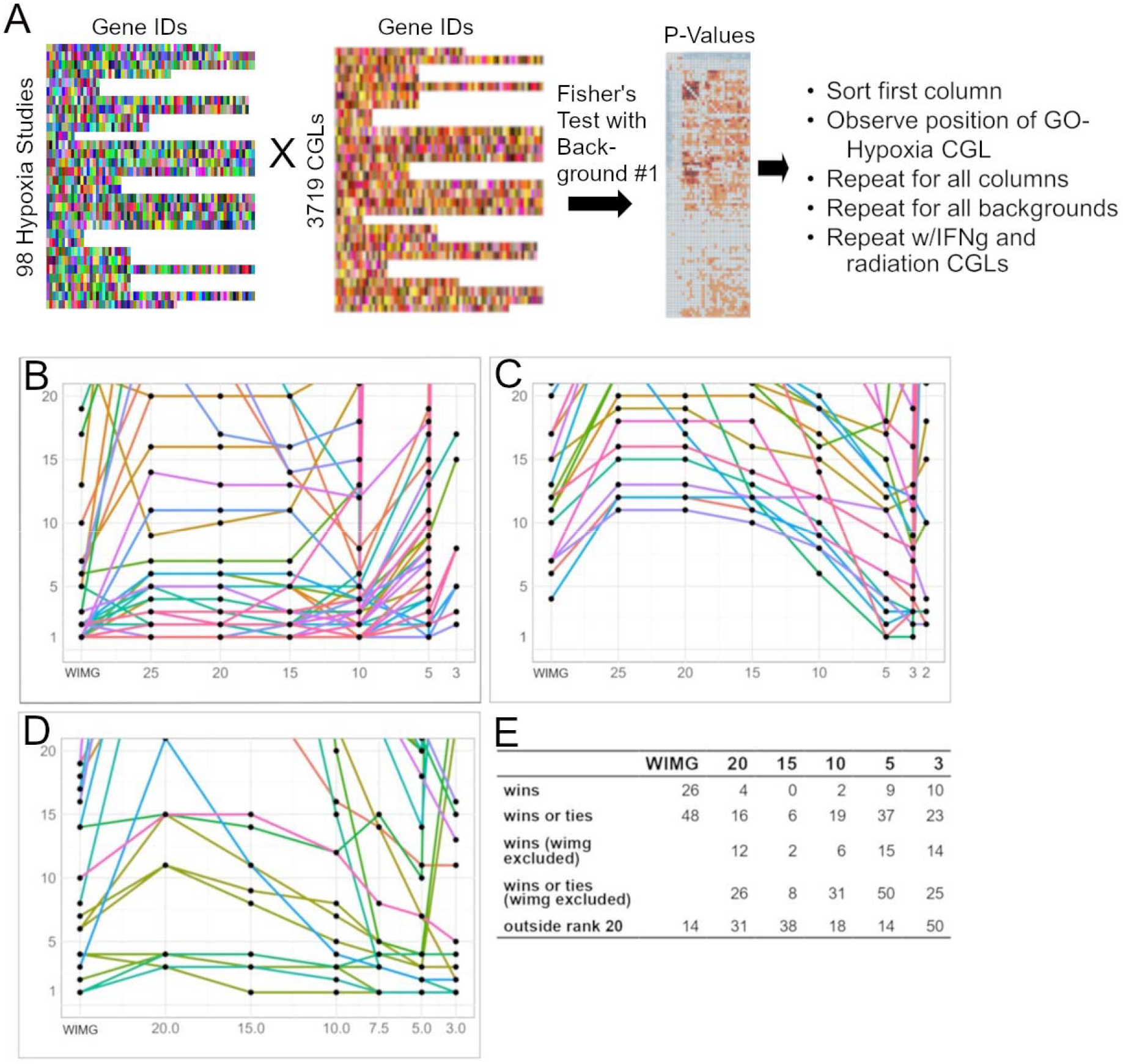
Exploration of backgrounds at which three GO-CGLs best match relevant lab-based studies. A. Scheme: Gene lists from 98 lab-based hypoxia studies are crossed with 3719 CGLs, generating 364,462 P-values. The rank of a GO-CGL representing hypoxia is noted against all 98 studies. The test is repeated for 6 backgrounds. The scheme is repeated for studies involving IFNg and radiation. B-D. colored curves represent individual lab-generated studies. The horizontal axis represents background divided by 1000 and the vertical axis represents ranking; a ranking of 1 means that the GO-CGL of interest out-competed 3718 other GO-CGLs. Rankings above 20 are not shown. B. Hypoxia studies. C. IFN-gamma studies. D. Radiation studies. E. Table shows combined performance of the three tests at 6 different background choices. A “win” means that the given background outperformed the other five for a particular study. Results including and excluding WIMG assigned backgrounds are both given. “Outside rank 20” refers to backgrounds that caused the GO-CGL to drop out of top-20 ranking for particular studies.

We repeated the above benchmarking procedure, this time examining the response to interferon gamma (GO:0034341 Response to Type II interferon) against 31 studies in which cells were treated with ifn-gamma (Fig. 3C). Remarkably, a background of 3000 performed best in driving the CGL to its highest rank, while backgrounds of 25000 and 20000 never drove the CGL to its highest rank. Rather, the two highest backgrounds caused the CGL to fall out of the top 20 rankings on 11 and 10 occasions.

A final benchmarking experiment involved the response to radiation (Fig. 3D). Here, we intersected 143 radiation-related studies with “GO:030330 DNA Damage Response Signaling Transduction By P53 Class Mediator.” The relevance of this CGL may not be obvious; nevertheless, a ranking of 20 or less was obtained on 24 occasions and rank #1 was seen on four occasions. Once again, low backgrounds (3000 and 5000) performed best in driving the CGL to its highest ranking versus other backgrounds, while backgrounds of 20000, 15000, and 10000 caused the CGL to fall out of the range of noticeability on 14, 13, and 10 occasions.

It is notable that the assigned backgrounds for the above CGLs, 12014, 6388, and 13722 respectively, are considerably lower than CGL backgrounds based on the size of the genome.

Figure 3E summarizes the above tests. WIMG assigned backgrounds were clearly optimal in causing the three CGLs to rise to their expected high rankings. In the absence of WIMG backgrounds, it is clear that low backgrounds performed optimally in driving CGLs to high rankings. Given the paucity of CGLs that are appropriate for the above procedure, we refrain from proposing that low CGL backgrounds would be universally optimal for all ESs. We would propose, however, that a curious researcher perform CGL/ES analysis under a variety of CGL backgrounds, even if *P*-values, which tend towards insignificance as backgrounds drop, do not impress.

Rankings of all studies at specified backgrounds are found in table S10.

### Study/study comparisons outperform study/GO-CGL comparisons

At the time of writing, the abridged WIMG database contains 70300 gene lists, of which 2604 are GO CGLs suitable for the following analysis. We performed Fisher’s exact test using the 124 most recently entered ESs against the 70300 gene lists (8.7*10^6^ *P*-values), observing the rankings of the best performing GO CGLs against each of the 124 lists (Table S4). Backgrounds were set at 20,000 for all CGLs. Fig. 4A illustrates the general algorithm. In all cases, an ES could be found that outranked (i.e. generated a more significant *P*-value than) the highest-ranking GO-CGL. Over the 124 tests, the best performing GO-CGL appeared as high as the 3^rd^ position, but as low as the 2268^th^ position. The average position of the highest ranked GO-CGL over the 124 tests was 468; i.e. typically, 467 other lists outperformed the top GO-CGL. Given the 2604 lists, one would naively assign a 50% probability of finding at least one GO-CGL within the top 27 ranks for each of the 124 tests, but this occurred in only 19 of the 124 cases (*P*= 5.4*10^-16^, binomial distribution).

**Figure 4.**
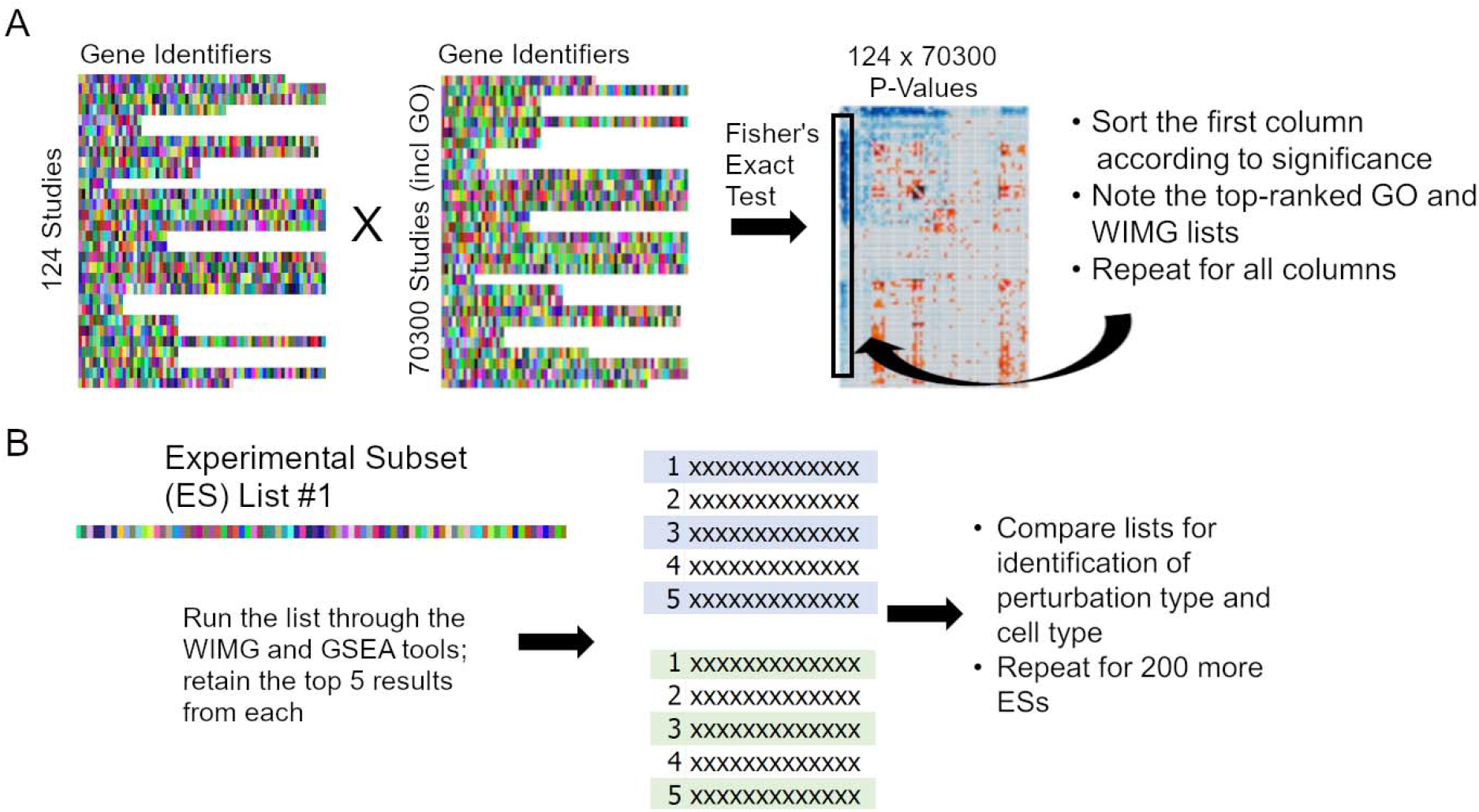
A. The scheme for ascertaining the extent to which CGLs and WIMG lists attain high-significance rankings relative to ESs. 124 RNA-seq studies were randomly chosen and crossed with 70,300 other studies (CGLs, WGCNA lists, WIMG lists, and a large number of ESs). For each of the 124 columns of *P*-values, the top ranking CGL and WIMG list is noted. B. Scheme for assessing WIMG’s competence in predicting cell type and perturbation type, given a random ES. A gene list is processed through both the GSEA and WIMG tools, the cell and perturbation types of the top five outputs are noted, the process is repeated for 200 more ESs, and results are compared.

Because of the parent-child structure of GO-CGLs, redundancy between GO-CGLs is expected. One could thus argue that it is unreasonable to expect typical GO-CGLs to breach the 27^th^ rank in our test. It could also be argued that since CGLs capture general processes over various species, cell types, and experimental conditions, it should not be surprising that very specific ES/ES combinations can be found that outrank CGL/ES combinations. Below, we show that CGLs can indeed perform strongly against ES/ES competition.

## WIMG lists

Currently, the WIMG database includes 265 “WIMG Lists.” These are created by three methods. First, studies representative of a well-defined process (e.g. hypoxia) may be gathered. Genes strongly altered throughout these studies are gathered into up- and down-regulation lists. These genes must be derived from unabridged datasets or supplemental tables, not via text-mining or scanning of papers. The second method involves clustering. As an example, *P*-values representing all disease-related ES/ES combinations may be generated. Clustering is then performed on a resulting disease/disease matrix that would currently contain 3*10^7^ *P*-values. Genes most representative of a particular cluster are accumulated into a list. The third method simply labels specific ESs as “WIMG exemplars.” For example, we note that genes down-regulated upon Raver2 knockdown in one study[30] repeatedly generate extreme *P*-values when compared against numerous other ESs. This process is not captured by GO-CGLs or by the other two classes of WIMG lists. We have labeled 24 ESs as “WIMG exemplars.”

We repeated the above test of 124 ESs against 70300 gene lists, this time observing the rankings of the 124 best-performing WIMG lists. Here, a WIMG list received top ranking in 10 of the 124 tests. The lowest ranking was 907, with an average ranking of 60. Given the 265 lists, one would expect a WIMG list to be found within the top 265.3 rankings in half of the 124 studies. However, WIMG lists were found in this region in 118 cases (*P*= 7.7*10^-24^, binomial distribution). Ignoring expectation values, the top ranked WIMG list still outranked the top-ranked GO-CGL in 100 of the 124 tests. We would argue that even over a mere 124 ESs, certain processes recur, and these are well-captured in WIMG lists.

The WIMG database contains 160 lists based on WGCNA clustering[31]. Typically, studies that utilize this procedure involve single-cell analysis, requiring large matrices to generate reliable gene-gene coexpression patterns. One may thus not expect such lists to overlap strongly with, for example, an ES involving the application of a cancer drug to a cell line. Despite this disadvantage, we repeated the 124 ES test against WGCNAs. Here, the highest ranking was 2, and the lowest 2044, with an average ranking of 558. WGCNAs breached the ranking expectation value on 72 occasions (*P*=.01, binomial test). As with most WIMG lists, the WGCNA method extracts patterns from raw data, as opposed to requiring that all genes within a list conform to easily-defined categories. This may explain WGCNA’s relatively strong performance compared to GO-CGLs.

For clarity, all WIMG lists were generated prior to entry of the 124 ESs into the database. The composition of all WIMG lists is available in Table S5. The results of the above GO/WGCNA/WIMG comparison are available in Table S4.

WIMG users may over-ride the backgrounds that WIMG attaches to GO-CGLs (see supplemental WIMG manual). When we repeated the GO/WGCNA/WIMG comparison tests with background-adjusted GO-CGLs, WIMG lists, and WGCNA lists, all tended to fare considerably worse against ES competition versus the non-adjusted (background = 20000) lists used in our initial tests. Nevertheless, background-adjusted WIMG lists outperformed expectations on 99 of 124 occasions, while background-adjusted GO-CGLs outperformed on only 8 occasions. In Table S4, the single most significant *P*-value derived from each of the 124 ESs against the remainder of the database is shown. One may term ESs associated with extreme significance as particularly “relevant” (conversely, ESs with weak maximal significance would be similar to ESs generated simply by scrambling gene lists). Interestingly, background-adjusted WIMG list rankings correlated with these maximal *P*-values (*P*= .013, t-test), which was not the case for non-adjusted lists. Following background-adjustment, WIMG lists correlate more strongly with the “relevance” of studies.

We considered the possibility that our typical sorting method for input ES data, that of dividing a gene’s log(fold-change) by the significance of its alteration, could alter the rankings of GO-CGLs vs ESs. We thus examined the results of deriving ES lists by first excluding all genes with significance >.05, and then sorting the remaining genes according to fold-change in a new test involving 132 ES input lists. In both cases, background adjustment was utilized for GO-CGLs, WGCNA lists, and WIMG lists. However, the sorting procedure made little difference in rankings. Over the 132 tests, GO-CGLs outperformed expectations on only four occasions using WIMG-typical sorting; the figure increases to five using the second sorting approach. WIMG lists outperformed expectations on 123 occasions using the first approach, and 121 times using the second. It should be noted that in these tests, WGCNA lists failed to outperform expectations. The results of these tests are available in Table S4 sheet “test2.”

### WIMG study/study comparisons are particularly informative

Despite a plethora of gene enrichment databases[32], few databases contain extensive ES data from which ES/ES comparisons may be drawn. Enrichr[33] and GSEA are two exceptions. We set out to compare the ability of WIMG, Enrichr, and GSEA to “predict” two known features of the input ES; the cell type and the perturbation type (knockout, disease, cell type, etc.). Enrichr was excluded from this test because of the absence of a single output list of top-ranked studies. 201 differential expression gene lists derived from studies appearing in the GEO database in December 2021 were generated. Each list, containing no more than 250 genes, was entered into the GSEA and WIMG tools, and the top ranking 5 output ESs or CGLs were examined for the presence of the cell type and perturbation type associated with the input. Fig. 4B illustrates the general algorithm for this test. In terms of perturbation, WIMG made correct predictions 25.2% of the time, while GSEA scored at 8.6% (*P*= 1.02*10^-11^, t-test, unequal variance). For cell type, WIMG scored at 35% and GSEA at 13.9% (*P*= 7.5*10^-12^) (see Table S6 and Methods).

The GSEA/WIMG comparison could be critiqued for a number of reasons. In particular, the composition of the December 2021 GEO database may not mirror the composition of the GSEA database, which certainly contains a larger ratio of CGLs to ESs than the WIMG database. We would point readers to another result involving the WIMG “keyword” feature. Each WIMG study is tagged with multiple keywords. Some of the keywords apply to many studies (e.g. brain, blood, disease), while others apply to a small fraction of studies (parasite, hdac, UPR). The appearance of these keywords over the top 500 ranked studies is compared to their appearance over the remainder of the database using a simple binomial test. The tool is unique to WIMG, so comparisons to other tools cannot be made. Table 1 offers some examples of the concordance between input data and output results.

**Table 1:** WIMG output (columns 2 and 3) strongly mirrors study input (column 1). A binomial test links keywords to each of the 201 studies found in Table S6. A subset of this table is shown above. Particularly significant keywords are listed in column 2. Column 3 highlights further observations regarding the top 5 ranked studies versus the remaining 70295 studies.

| STUDY | WIMG keywords:<br>negative log(p)>10 | NOTES |
| --- | --- | --- |
| upregulated in ovcar3 line on boris oe vs kd (GSE166767) | female, epithelial, line, ovary, adherent | 5/5 WIMG studies involve ovcar3 |
| downregulated in a549 line on il17d oe (GSE191111) | carcinoma, lung, epithelial, line, adherent, male, female, breast | 5/5 WIMG studies involve a549 cell line |
| enriched on dmd clip (vs input) (GSE128318) | rip, epithelial, adherent, line | note the correct prediction of an RNA-IP experiment |
| upregulated in gingival epithelial cells on irf1 kd (both w/hsv infection) (GSE184463) | line, adherent, epithelial, kd | WIMG predicted a knockdown in 5/5 studies |
| upregulated in mouse follicular B cells on cic ko (GSE163455) | lymphocyte, b-cell, blood | WIMG 4th ranked study involves cic chipseq |
| downregulated in tregs on stimulation w/anti-tnfr2 agonistic antibody vs no stimulation (GSE161835) | lymphocyte, blood, treatment | 4/5 WIMG studies involve t-cell activation |
| upregulated in ipsc-derived astrocytes from schizophrenia-affected twin vs unaffected (GSE191248) | stem, neural, fibroblast | 2/5 WIMG studies involve astrocytes |
| upregulated in mouse e13.5 heart on prdm16 ko (GSE179386) | heart, mouse, cell type, embryo, brain, ko, neuron | 2/5 WIMG studies involve mouse embryonic heart |
| downregulated in mouse cd11c+ cells on high cholesterol diet vs control (Apolipoprotein E derived from CD11c+ cells ameliorates atherosclerosis) | blood, mouse, lymphocyte | WIMG dendritic keyword -log(p) = 6.47 |
| upregulated in mouse aml line on bcl11a oe (both w/trib1 oe) (GSE147797) | Blood, mouse, lymphocyte, bone marrow, ko, macrophage | WIMG leukemia keyword -log(p) = 7 |
| upregulated in ovary-metastatic tissue vs primary gastric adenocarcinoma (GSE191139) | cell type, brain, nerve, ovary | WIMG cancer tissue keyword -log(p) = 8 |
| downregulated in ovary-metastatic tissue vs primary gastric adenocarcinoma (GSE191139) | disease, cancer tissue, stomach, blood | WIMG metastasis keyword -log(p) = 6.98 |
| upregulated in huvecs on shp2 kd (GSE191056) | endothelial, umbilical, ifn | 4/5 WIMG studies involve endothelial cells |
| upregulated in hnecs on 14d hrv16 infection (GSE190344) | ifn, cytokine, virus, RNA virus, infection, treatment, umbilical | WIMG 5th ranked study involves nasal tissue upon rhinovirus infection |
| downregulated in CD11c+ cells from 9m vs 5d mouse brain (GSE190713) | mouse, ko, hippocampus, brain | 2/5 WIMG studies involve Alzheimer's |
| downregulated in mouse follicular B cells on cic ko (GSE163455) | Blood, lymphocyte, b-cell, mouse, ko, spleen | 4/5 WIMG studies involve B-cells |
| downregulated in mouse cerebellum on aptx ko (GSE174841) | brain, mouse, cerebellum, ko, cortex | note that cerebellum is a significant keyword |
| downregulated in gingival epithelial cells on irf1 kd (both w/hsv infection) (GSE184463) | ifn, cytokine, breast, epithelial, treatment, gland, line | WIMG study #5 involves irf1 |
| downregulated in cd138+ multiple myeloma cells at relapse vs diagnosis (GSE162403) | blood, cell type, lymphocyte, pbmc | WIMG study #1 relates to CML; study #3 to AML |
| upregulated in keloid tissue vs healthy (GSE190626) | cancer_tissue, fibroblast, disease | WIMG #1 ranked study involves keloid tissue |
| downregulated in rat striatum with (vs w/o) physical exercise following stroke (GSE190355) | brain, hippocampus, mouse, rat, ko, striatum | note WIMG keyword "rat"; note WIMG keyword "striatum" |
| upregulated in mescs on ubn2 kd (GSE177026) | embryo, stem, mouse, muscle, ko | 5/5 WIMG studies involve mescs |
| upregulated in squamous vs non-squamous lung tumors (GSE190265) | skin, breast, cancer_tissue, keratinocyte, esophagus, lung, gland, head/neck, bladder, tongue | 5/5 WIMG studies involve lung squamous cell carcinoma |
| upregulated in mouse MLL-AF9 leukemia stem cells on tifab oe (GSE178853) | blood, leukemia | note the "leukemia" keyword; 4/5 WIMG studies relate to leukemia |
| upregulated in HO-1-N-1 line on C. parapsilosis treatment vs heat-killed C. parapsilosis treatment (GSE169278) | epithelial, adherent, female, line | top ranked WIMG study involves fungus (aspergillus) infection |
| downregulated in patient-derived rhabdomyosarcoma lines on ehmt1 kd (GSE179049) | female, line | 4/5 WIMG studies relate to sarcoma |
| upregulated in mouse b-cells heterozygous for tbk1 (GSE188659) | blood, lymphocyte, mouse, b-cell | 2/5 WIMG studies involve mutation; 1 involves haploinsufficiency |
| upregulated in mouse microglia on tamoxifen-induced Cre activity (vs no induction) (GSE190207) | ifn, cytokine, virus, RNA virus, infection, treatment, macrophage, mouse, microglia | note the WIMG microglia keyword |

We would ask readers to peruse Table S6 and enter their own ESs into the WIMG “Fisher” tool for further evidence of the ability to predict the nature of the input set. Researchers, of course, are not interested in learning that their pancreatic adenocarcinoma study is indeed a cancer study. However, WIMG’s ability to correctly mirror *known* aspects of the input to users should enhance confidence in its ability to properly reflect back *unknown* or unexpected aspects of the study. If a knockout results in output that mirrors the trends seen in glioma, or an antiretroviral treatment reverses Alzheimer’s trends, or an exercise regimen reverses trends seen in aging studies, researchers may wish to take note.

### Tackling gene set overlap

After submitting an ES for analysis, the CGL output list may, depending on the tool, contain hundreds or even thousands of ranked CGLs. Unfortunately, this does not mean hundreds of insights have been generated, as highly ranked outputs tend to be quite similar in content. This is the gene set overlap problem. A number of solutions have been employed[7, 34–36]. However, gene set overlap may still remain strong. For example, utilizing one of these solutions, we enter a list of genes upregulated in nasal epithelial cells on ifn-lambda treatment[37]. Selecting a redundancy reduction feature, the first and second ranked outputs, involving the response to type I interferon and the response to gamma interferon, still contain 16 genes in common. Applying Fisher’s exact test to these two sets, we derive *P* < 10^-100^.

WIMG takes an extreme approach to eliminating redundancy. Using the site’s “Third Set” tool, the user enters two gene sets that are known to intersect. The tool seeks a third gene set that overlaps with the first, “central” set, but does not overlap significantly with the second set (i.e. *P*>.05). Inputting the ifn-lambda experimental data and genes found in GO:0034340 (response to type I interferon), and setting the “experiment” filter to “external list”, which eliminates all ESs from consideration, the tool is actually unable to find a second GO-CGL that intersects significantly with the central set but not the secondary set. Removing the “external list” restriction, however, the tool is indeed able to find numerous studies that overlap with the ifn-lambda study but not the GO-CGL. The second-ranked study involves genes upregulated in the hela line on ifn-gamma vs ifn-beta (a type I interferon) treatment[38] (Fig 5B). This result “makes sense”, perhaps heightening confidence that the top-ranked study, involving a novel αVβ3 inhibitor, relates specifically to ifn-gamma metabolism[39].

**Figure 5.**
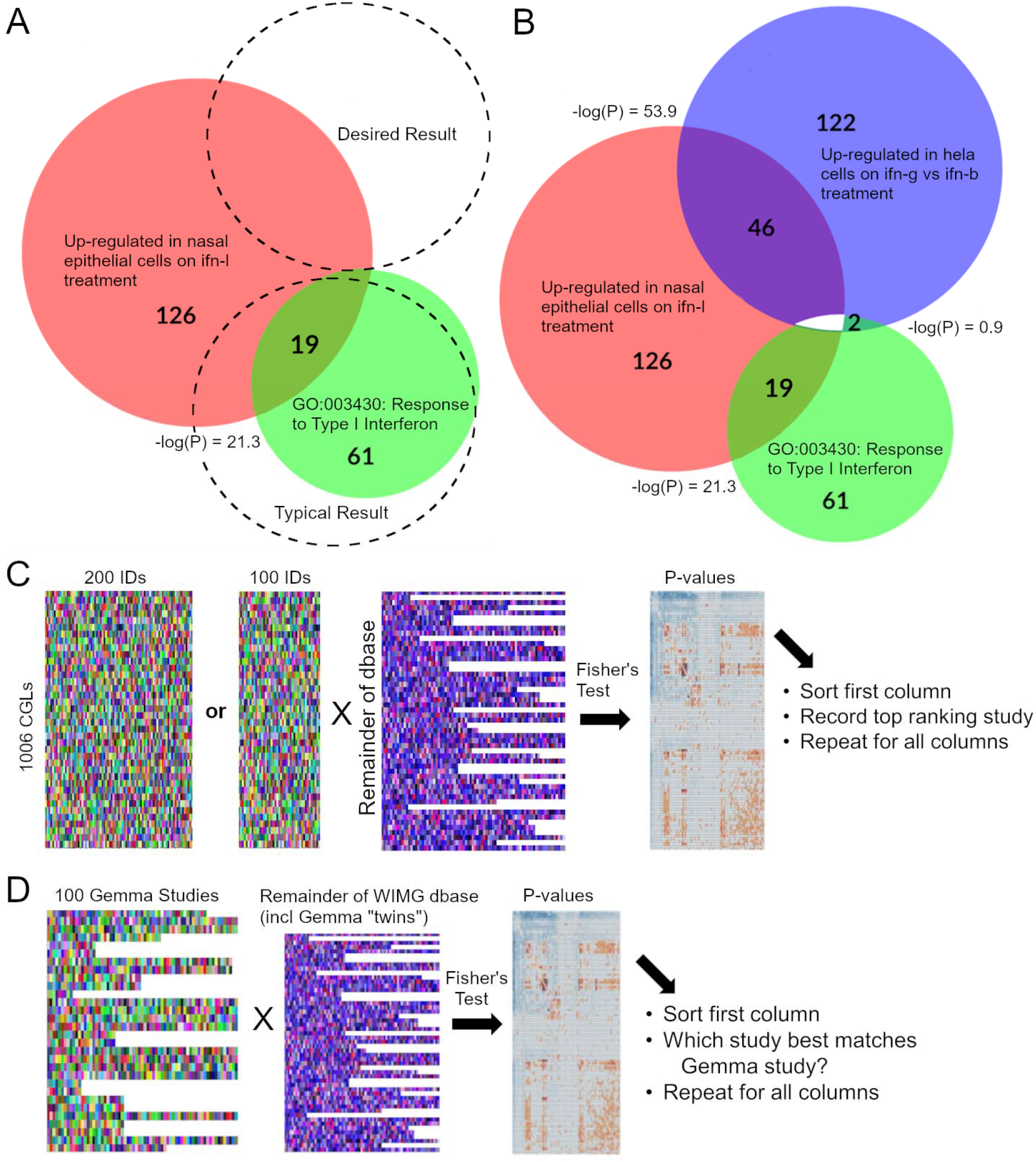
Tackling redundancy of output sets, the cut-off problem, and batch control. A. A user submits a 126-member list to a gene enrichment tool. The top ranking CGL is returned (green). Further insight into the nature of the submitted list may be hindered because other high-ranking CGL outputs tend to be similar to the top-ranked CGL. B. WIMG redundancy reduction. The central IFN-lambda gene set and the secondary GO:0034340 gene set were entered into the “Third Set” tool. The tool found a third gene set that intersects strongly with the central set, but not the second set. *P*-values for each of the three intersections are given. C. Scheme for testing gene set length cut-offs. After sorting genes in 1006 CGLs according to frequency of perturbation, all CGLs are truncated to 200 and 100 member lengths. The two sets of CGLs are both crossed with the complete WIMG database, a *P*-value matrix is generated, and the ability of the two sets to draw CGLs to top-ranking positions is evaluated. D. Assessing the benefits of batch control at the level of gene enrichment. 100 Gemma lists are crossed with the entire WIMG database, which already contains the 100 studies (with no additional batch control) on which Gemma batch control was conducted. The question to be resolved is whether or not the unadjusted Gemma “twins” rank as the best matches to the Gemma lists.

The general inability to find CGLs that do not themselves overlap, but do overlap with a central ES, may have necessitated the construction of redundancy reduction algorithms that cannot entirely eliminate significant intersection between output sets. When ESs are allowed as outputs, however, a thoroughly restrictive approach may produce results.

Note that the tool has utility beyond redundancy reduction. If a user has reason to believe that a dominant set of genes is “blocking” an otherwise interesting result, the third set tool effectively removes the dominant group, as well as gene lists that contain content similar to that of the dominant group. The blocking set could range from a specific list of genes typically associated with innate immunity, which could be hiding subtleties involved in the response to a particular virus, to a more general list of genes which evince disproportionate differential expression over a large range of phenotypes[40].

One important drawback to the WIMG approach is the fact that two sets (as opposed to one set) must first be identified for input, meaning that the “Third Set” approach may serve as a complement to existing tools.

### The cut-off problem

Two basic algorithms are most commonly employed for gene set analysis; over-representation analysis (ORA) and functional class scoring (FCS)[35, 41]. WIMG uses an ORA-based algorithm. When discussing the weaknesses of ORA against FCS, it is commonly pointed-out that ORA requires input of a subset of all identified genes, whereas FCS requires only that all genes be ranked according to some criteria. The truncation of ORA input lists may be considered somewhat artificial; genes differentially expressed at *P* = .049 may make the cut, but genes with *P*= .051 are ignored. A related issue is the fact that ORA tools treat each gene equally; genes differentially expressed with extreme significance are no more valuable than genes that barely make the cut.

To some extent, WIMG resolves the above critiques. This is because 1) 81% of the gene lists in the WIMG database are sorted and 2) users may elect to shorten a WIMG gene list such that only the most significantly altered genes remain. When large numbers of genes show statistically significant differential expression, lists are sorted according to log(fold-change) divided by significance. Otherwise, lists are sorted by log(fold-change) alone. Users can examine both full and truncated WIMG lists, looking for *P*-value alterations as the list shortens. If, for example, the most significant *P*-value is found against a truncated version of a WIMG list, one can conclude the excluded genes are fairly irrelevant. In the case of FCS, a “leading edge” subset of the complete input list is generated. As with any ORA tool, the WIMG user is free to alter the length of an input list. However, with WIMG, database lists may be truncated as well, reflecting the view that input lists and database lists should be treated equivalently.

We intersected 1,006 WIMG ESs and CGLs, all pruned to exactly 200 genes, against the remainder of the WIMG database, generating 7*10^7^ *P*-values. For each of the 1,006 lists, the single most significant *P*-value against the remainder of the database was recorded. We then repeated the same exercise, this time pruning the 1,006 lists to 100 genes each. We refer to this as the “halving test” (Fig 5C). A large, “well-focused” list of genes should always outperform a short, equally focused list in tests of significance, as increased intersection between an input and a database set will overwhelm minor alterations in the number of genes in these two sets. Nevertheless, in 6.8% of cases significance increased when the 1,006 lists were halved in size, suggesting that the remaining 100 genes in the 200-member set are of minimal importance, at least with respect to whatever insights can be gleaned from the two gene lists in question.

Most WIMG GO-CGLs are sorted according to “perturbability.” In brief, a frequency table of genes appearing over all WIMG ESs that involve a perturbation (e.g. a disease or knockout) is generated. Currently, EGR1 is the most commonly perturbed gene in the database, appearing in 7% of ESs. Genes in GO-CGLs are then sorted according to these frequencies. If EGR1 is present in a GO-CGL, it will rank first in the list. Housekeeping genes, while common and critical to many of the processes that GO-CGLs seek to embody, receive the lowest rankings. Just as housekeeping genes and other rarely altered genes tend not to be found in lab-based ESs, they may also be excluded from WIMG GO-CGLs via truncation.

We repeated the halving test, this time focusing on 1,004 sorted GO-CGLs with at least 200 members (which were subsequently trimmed to precisely 200 members). GO-CGLs were not allowed to intersect with GO-CGLs, but were allowed to intersect with WIMG-CGLs. In this case, significance increased 34% of the time when GO-CGLs were halved. The pruning of GO-CGLs may actually improve their relevance to ESs.

WIMG does not offer an option to simultaneously test an input set against multiple database gene set lengths, due simply to the processing required. Currently, one would need to generate an initial ranked output list with the “Fisher” tool, trim the database via the “Restrict IDs” option, generate a second ranked list, and compare the two lists.

### Batch Control

Despite efforts to strictly follow experimental protocols, surprisingly large effects on transcript or protein counts in test and control samples may be generated by minor or unobserved variations in experimental timing, location, reagents, and technique. A number of methods have been devised to adjust data to account for such batch effects[42–44].

Currently, only 0.5% of gene lists within the WIMG database have been derived via workflows that include batch control. These lists are obtained from the GEMMA database[45], which reprocesses GEO datasets to account for possible batch effects. We compared 100 RNA-seq-based gene lists already found within the WIMG database to the parallel batch-controlled lists generated from the GEMMA database (Table S10). The studies were required to have a GEMMA quality index of 0.3, indicating that batch effects were indeed detected between samples. We also required that covariates be minimized, as this is already WIMG practice (e.g. analysis of the effect of a drug on a single cell line is preferred over analysis over multiple cell lines). Fig. 5D illustrates the algorithm for analysis. Using Fisher’s exact test for the comparison, the median –log(*P*) was 110, with 13 comparisons giving a value greater than 300, and a low value of 0.78. Perhaps more importantly, in all but 12 cases the *P*-value generated by the above GEMMA/WIMG comparison was the single most significant result against *P*-values generated via comparisons of GEMMA data against 77,300 other gene sets in the WIMG database. Thus, while fold-changes and significances of individual genes of interest may fluctuate after batch adjustment, the absence of batch control does not generally appear to have important effects at the level of comparisons between gene sets. Nevertheless, we anticipate increasing the percentage of gene lists generated via batch control workflows in the WIMG database.

## Discussion

Cursory experimentation reveals that the *P*-values generated in gene enrichment, as well as the ensuing ranking of various gene lists, can be quite sensitive to assumptions regarding the size of gene set backgrounds. Merely summing citations of four of the most popular enrichment tools/algorithms[18, 22, 46, 47], we derive a figure of 180,000, implying the usage of these tools in approximately the same number of papers. Such a figure does not include numerous “minor” tools, commercial platforms (e.g. IPA), failures to cite, or the occasions in which experimentalists use such tools to drive hypotheses without publication. Not only are large sets of differentially regulated genes cognitively intractable without some form of summarization, meaning that it is difficult to perform transcriptomics or proteomics without enrichment, but some form of enrichment is also considered a best practice[48]. Given the critical role of gene enrichment in modern biology, it would seem surprising that the optimization of background definitions has received scant attention relative to the algorithms themselves.

We evaluated various approaches to gene set backgrounding by means of Shannon information. These metrics are suggested as a tool in the assessment of gene enrichment approaches. Imputed backgrounds performed strongly against standards. The results are not without nuance, however. In one case, merely setting CGL and ES backgrounds to 20,000 led to highest entropy; this case was also characterized by anomalous skew. Increases in information resolution must therefore be considered alongside other factors, including skew, kurtosis, and indicators of biological “reality.”

When comparing our evaluations of CGL/ES and ES/ES *P*-value matrices, it should be pointed out that the majority of CGLs represent generalized biological processes (e.g., “cell cycle”) and are not resolved by tissue type. Consequently, they are best modeled by a lower background that reflects average detectability across the proteome or transcriptome. In contrast, imputation for experimentally derived gene lists utilizes tissue-specific abundance data, allowing for more precise calibration of the background. If CGLs were simply coupled with recommended tissue-specific backgrounds, the gap between generalized ontology and experimental reality would likely narrow.

Another evaluation of backgrounding was conducted at the level of specific GO-CGLs, with the goal of finding the backgrounds that best drive these CGLs to their expected high ranks versus irrelevant CGLs. Here, imputed backgrounds were clearly optimal; merely setting all CGLs to a single background, from 3,000 to 25,000, consistently underperformed versus the approach of rational background estimation. In the absence of imputation, low backgrounds strongly outperformed high backgrounds. This result amplifies the finding in the entropy analysis wherein simply setting CGL backgrounds to 5,000 outperformed the 10,000 and 20,000 settings.

We have shown that GO-CGLs, and presumably other similarly constructed lists, perform poorly when ranked against ES competition. This is not simply because “averaged-out” exemplars of particular processes cannot be expected to fare well against very specific ES/ES combinations, as WIMG lists outperformed ES competition on numerous occasions. WIMG evinces a strong ability to reflect back known aspects of input data to users. Put differently, the tool maximizes true positive identifications. WIMG also tackles standard enrichment obstacles such as gene-set overlap and the problem of arbitrary cutoffs in ORA algorithms. We suggest the sorting of genes in CGLs and the assignment of backgrounds to CGLs as means to improve their relevance to real-life experimental subsets.

Given an input list of genes, the ability to predict the cell type and, to a lesser extent, perturbation type, may come as a surprise to some. For example, one might assume that differentially expressed genes (DEGs) in a p53 knockout experiment in the a549 cell line would tend to best overlap with DEGs in other studies involving perturbation of p53 or its pathway. Instead, it is more likely that the study will simply overlap with other a549 studies. Repeatedly, we see that cell line experiments can be separated from in vivo studies, drug studies are more likely to overlap with other drug studies (as opposed to, say, knockout studies), disease studies tend to best match other disease studies, etc. These observations have practical consequences as, in the course of examining studies for addition to the database, we routinely see datasets in which a drug is applied to multiple cell types, but each cell type is represented by only one test and control sample. The result is typically a list of genes without any significant differential expression, which may not be recoverable with batch correction.

There appears to be tension between general, easily understood, yet relatively insignificant categories represented by CGL outputs and specific, sometimes obscure, significant ES outputs. If, for example, we found that a list of genes neighboring ultraconserved DNA stretches[49] overlapped strongly with those involved in oxidative phosphorylation, hints of a hypothesis may be seen: these stretches are involved in the preservation of ancient, critical metabolic genes. In truth, this is not the case; instead, we find that ultraconserved genes overlap the genes targeted by the transcription factors pcgf2, phc1, and jarid2[50]. It is perhaps more difficult to formulate a hypothesis in this case, though a researcher whose work involves these transcription factors may be able to draw a link. We may also imagine a case where ultraconserved stretches indeed neighbored oxidative phosphorylation genes, but at far lower significance than the overlap with the targets of particular transcription factors. In the absence of ES data, the researcher may prioritize CGLs whose appearance is actually secondary to the actual mechanistic drivers.

As with any gene enrichment tool, WIMG should not be considered a one-stop solution. We have already pointed out some features that may suffer in comparison to other tools. To these, we add the fact that our emphasis on database bulk may come at the cost of quality control and speed of output. Major sources of data include the GEO and GREIN[51] databases. In most cases, no batch correction of test vs control sets beyond that offered on these sites is conducted. Another source is supplemental datasets, which we take at “face value.” Datasets with obvious gender bias (e.g. an excess of males in a test group vs a control group), unusually small backgrounds, and targeted arrays (e.g. those focused only on cancer genes) are generally excluded from the database, however. Our assumption is that noisy, information-poor sets in the database generally will not percolate to the top output ranks; this could be proven false.

Another concern involves benchmarking. A number of attempts to compare performances of various gene enrichment tools have been made[52–54]. In all cases of which we are aware, however, CGLs are used as “gold standards” for analysis; a tool that best matches a leukemia ES to a cancer CGL is declared winner. Even synthetic benchmarking approaches, which utilize simulated data, ultimately rely on CGLs as the “ground truth” target. Given our numerous concerns with the usage of CGLs, we refrain from this particular exercise, opting instead for “within house” benchmarking tests (e.g. our estimates of *P*-matrix information content and tests of the ability of particular backgrounds to draw CGLs to their expected high ranking gene enrichment output positions).

WIMG can be classed as an ORA approach. The question arises of whether FCS overcomes the above concerns regarding CGLs with undefined backgrounds. Briefly, let us assume a cancer ES with background of 20000 and a cancer CGL with an effective background of 4000. One would then expect that 80% of truly relevant cancer genes could be absent in the CGL; such true positive genes actually subtract from the FCS running sum. There is no reason, however, that FCS algorithms cannot be altered to accommodate CGLs and ESs with unequal backgrounds.

Future directions for WIMG include a more dynamic interface with the ability to generate relevant graphics. Ideally, the run-time of some tools should decrease. New features may include an option to analyze input data via an FCS approach. Most importantly, we believe that the sheer volume of ESs in the WIMG database creates an opportunity for AI-based analysis of input ESs and the database as a whole.

WIMG tools may be accessed at WhatIsMyGene.com. The site is funded by donations.

## Methods

10532 GO-CGLs were downloaded from gsea-msigdb.org[18] . Gene sets with fewer than 32 or more than 1000 annotations were then excluded. All sets were then sorted according to the frequency with which genes are perturbed within the WIMG database. Each GO-CGL was assigned an “abundance ranking” by first assigning each gene within the set a ranking based on the gene’s abundance within the whole human proteome or mouse brain proteome, and then calculating the average rank for all genes within the set. To avoid confusion, we should point out that the WIMG database actually contains 3803 external CGLs at the time of writing, but only the 2604 that have been sorted were used for downstream analysis. The GO list abundance rankings are available as Table S8.

It should be noted that the data used in fig. 1d is atypical in the sense that the authors included nonsignificant GO terms in their table (s4 “GOCC_UE-RAB3GAP1”). Had only significant GO terms been listed, the correlation between abundance and significance would have weakened. The exercise of correlating GO term significance with GO list abundance rankings can, of course, be performed for any input set of choice by utilizing any GO enrichment tools.

Background estimates (imputations) of gene lists can be derived by first locating a ranked abundance list of entities relevant to tissue, species, and molecule (protein or transcript). The gene list is then merged with the abundance list, allowing average abundance in the gene list to be calculated. This figure is then doubled to reflect the range of abundances in the gene lists, from most to least abundant. More sophisticated approaches can be considered.

The “reverse Fisher” method of background estimation is based on the observation that an estimate can be numerically derived by repeatedly solving Fisher’s exact test with three known parameters (sizes of input set A and B, as well as the size of intersection between A and B) and a range of background values, and then retaining the background at which *P* reaches a maximum (i.e. the least significant *P*). This procedure, however, can be simplified to an analytical approach wherein the size of the input list is multiplied by the size of a list of the top most abundant genes; this product is then divided by the size of the intersection. The analytical approach is currently utilized by WIMG.

A gene list is simply a set of 1 or more unique gene IDs. A gene list may be evaluated for similarity to other gene lists via Fisher’s exact test with four parameters as input: the length of list A, the length of list B, the size of the A|B intersection, and the joint background of the two lists. Given N gene lists, then, N^2^/2-N study/study overlaps and corresponding *P*-values may be generated. After reducing the 2-dimensional matrix of *P*s to a single list, standard statistics may be applied. We chose Shannon entropy because it is the most direct quantifier of information in such data; a single-peak distribution may have relatively high standard deviation and/or low kurtosis and nevertheless exhibit relatively low information content, as results still tend to concentrate in a single peak.

To quantify the discriminative power and information content of the *P*-value distributions generated by various background methods, we calculated Shannon Entropy (H). As our *P*-values are signed (negative for enrichment, positive for depletion), and their raw ranges can vary significantly across methods, we employed a standardized approach to ensure fair comparison. First, for each set of *P*-values, we performed a Z-score transformation. This rescaled each distribution to have a mean of zero and a standard deviation of one, removing the distorting influence of varying means and overall spread (scale). This step allows for a direct comparison of the fundamental shapes and complexities of the distributions, independent of their absolute numerical magnitudes. Next, these standardized scores were discretized into bins to approximate their probability density function. To maintain rigor, we used a fixed global binning strategy: a universal range of -10 to +10 standard deviations was divided into 100 equally spaced intervals. This created a uniform ‘ruler’ against which all standardized distributions were measured. The probability (p_i_) of a score falling into the i_th_ bin was then calculated as the count frequency divided by the total number of observations. Shannon Entropy was computed as the negative sum of p_i_ multiplied by the base-2 logarithm of p_i_ across all bins. This multi-step process—standardization followed by fixed-bin entropy calculation—eliminates the possibility of entropy scores being artificially inflated by wider dynamic ranges or automated bin-count variances inherent to specific methods. It ensures that differences in H truly reflect variations in the complexity and distinctness of the underlying statistical distributions, representing the genuine information content. Settings remained consistent for all datasets analyzed. To quantify the relative difference in information content between two approaches, we report the increase in discriminative power as 2^(H1 - H2), representing the proportional increase in the effective number of distinguishable states.

Data in the “Benchmarking CGL backgrounds” section was generated as follows. Gene lists for 3719 GO-CGLs were combined with gene lists for ESs of interest in a matrix. *P*-values for all CGL/ES intersections were generated at specified CGL backgrounds. Backgrounds for ESs were not altered from those found in the WIMG database. The ranking (according to significance) of the target CGL (e.g. Response to Oxygen Levels) was recorded for each ES. The procedure was repeated for all backgrounds.

To compare rankings of GO-CGLs, WGCNA clusters, and WIMG lists against input ESs, the following steps were taken. First, studies entered into the GEO database between October and December 2021 were selected. This relatively short time-frame is intended to minimize the possibility of “cherry picking” studies that would tend to prove our points. Individual input gene lists derived from these studies were then entered into the WIMG Fisher tool. The ranks of the single highest-ranking GO-CGL, WGCNA cluster, and WIMG list were recorded, and the exercise repeated with the next study. Expected top rankings for GO-CGLs were calculated simply by dividing the total number of gene lists in the WIMG database at the time of the test by the number of GO-CGLs in the WIMG database; the same applies for WGCNA clusters and WIMG lists. Test #1 was conducted over two background settings (all CGL backgrounds are fixed at 20000 and all CGL backgrounds are estimated using “reverse Fisher”). Test #2 was conducted using two methods of sorting input data (dividing log(fold-change) by significance and by excluded all insignificant genes and then sorting by fold-change).

Regarding the GSEA-WIMG comparison (Table S6), in the case of an experimental “cell type” designation, we required that a comparison between two cell types be made (as opposed to “liver cluster 17”).

For WIMG lists, clustering is performed using a matrix of Fisher *P*-values generated by intersecting relevant studies. Optimizing cluster numbers is performed with the R fviz_nbclust function from the factoextra package. Clustering is performed with Cluster 3.0[55] using k-Means with the Euclidean distance similarity metric. The resulting .kgg file contains the names of studies according to cluster. Gene IDs from within these studies are then tabulated. The most frequent genes in a cluster are compiled into a WIMG list and a background is assigned to the list.

Graphics for figures 1 and 2 were generated using the Matplotlib 3.7.1[56] and Seaborn 0.12.2[57] libraries executed in the Python 3.9 environment. The R package ggplot2[58] was used for line plots in figure 3.

## Competing Interests

The authors declare no competing interests.

## Funding

The study is not funded by any institution.

## Author’s Contributions

KH performed data collection and analysis, programming, and writing. TS performed programming, data analysis, and graphics generation.

## Notes

### Competing Interest Statement

The authors have declared no competing interest.

### Summary of Updates

We have improved our Shannon entropy analysis and have changed one figure relevant to this improvement. Minor edits have also been made throughout the paper.

## REFERENCES

1. The Gene Ontology Consortium, Aleksander SA, Balhoff J, et al (2023) The Gene Ontology knowledgebase in 2023. GENETICS 224:iyad031. 10.1093/genetics/iyad031

2. Kanehisa M, Furumichi M, Sato Y, et al (2023) KEGG for taxonomy-based analysis of pathways and genomes. Nucleic Acids Res 51:D587–D592. 10.1093/nar/gkac963

3. Thomas PD, Campbell MJ, Kejariwal A, et al (2003) PANTHER: a library of protein families and subfamilies indexed by function. Genome Res 13:2129–2141. 10.1101/gr.772403

4. Gillespie M, Jassal B, Stephan R, et al (2022) The reactome pathway knowledgebase 2022. Nucleic Acids Res 50:D687–D692. 10.1093/nar/gkab1028

5. Liberzon A, Birger C, Thorvaldsdóttir H, et al (2015) The Molecular Signatures Database (MSigDB) hallmark gene set collection. Cell Syst 1:417–425. 10.1016/j.cels.2015.12.004

6. Khatri P, Drăghici S (2005) Ontological analysis of gene expression data: current tools, limitations, and open problems. Bioinforma Oxf Engl 21:3587–3595. 10.1093/bioinformatics/bti565

7. Tarca AL, Draghici S, Bhatti G, Romero R (2012) Down-weighting overlapping genes improves gene set analysis. BMC Bioinformatics 13:136. 10.1186/1471-2105-13-136

8. Haynes WA, Tomczak A, Khatri P (2018) Gene annotation bias impedes biomedical research. Sci Rep 8:1362. 10.1038/s41598-018-19333-x

9. Tomczak A, Mortensen JM, Winnenburg R, et al (2018) Interpretation of biological experiments changes with evolution of the Gene Ontology and its annotations. Sci Rep 8:5115. 10.1038/s41598-018-23395-2

10. Davies MN, Meaburn EL, Schalkwyk LC (2010) Gene set enrichment; a problem of pathways. Brief Funct Genomics 9:385–390. 10.1093/bfgp/elq021

11. Gillis J, Pavlidis P (2011) The impact of multifunctional genes on “guilt by association” analysis. PloS One 6:e17258. 10.1371/journal.pone.0017258

12. Fulcher BD, Arnatkeviciute A, Fornito A (2021) Overcoming false-positive gene-category enrichment in the analysis of spatially resolved transcriptomic brain atlas data. Nat Commun 12:2669. 10.1038/s41467-021-22862-1

13. Tamayo P, Steinhardt G, Liberzon A, Mesirov JP (2016) The limitations of simple gene set enrichment analysis assuming gene independence. Stat Methods Med Res 25:472–487. 10.1177/0962280212460441

14. Rocha JJ, Jayaram SA, Stevens TJ, et al (2023) Functional unknomics: Systematic screening of conserved genes of unknown function. PLoS Biol 21:e3002222. 10.1371/journal.pbio.3002222

15. Edwards AM, Isserlin R, Bader GD, et al (2011) Too many roads not taken. Nature 470:163–165. 10.1038/470163a

16. Stoeger T, Gerlach M, Morimoto RI, Nunes Amaral LA (2018) Large-scale investigation of the reasons why potentially important genes are ignored. PLoS Biol 16:e2006643. 10.1371/journal.pbio.2006643

17. Gillis J, Pavlidis P (2012) “Guilt by association” is the exception rather than the rule in gene networks. PLoS Comput Biol 8:e1002444. 10.1371/journal.pcbi.1002444

18. Subramanian A, Tamayo P, Mootha VK, et al (2005) Gene set enrichment analysis: a knowledge-based approach for interpreting genome-wide expression profiles. Proc Natl Acad Sci U S A 102:15545– 15550. 10.1073/pnas.0506580102

19. Raudvere U, Kolberg L, Kuzmin I, et al (2019) g:Profiler: a web server for functional enrichment analysis and conversions of gene lists (2019 update). Nucleic Acids Res 47:W191–W198. 10.1093/nar/gkz369

20. Mi H, Muruganujan A, Ebert D, et al (2019) PANTHER version 14: more genomes, a new PANTHER GO-slim and improvements in enrichment analysis tools. Nucleic Acids Res 47:D419–D426. 10.1093/nar/gky1038

21. Timmons JA, Szkop KJ, Gallagher IJ (2015) Multiple sources of bias confound functional enrichment analysis of global -omics data. Genome Biol 16:186. 10.1186/s13059-015-0761-7

22. Huang DW, Sherman BT, Lempicki RA (2009) Systematic and integrative analysis of large gene lists using DAVID bioinformatics resources. Nat Protoc 4:44–57. 10.1038/nprot.2008.211

23. Zheng Q, Wang X-J (2008) GOEAST: a web-based software toolkit for Gene Ontology enrichment analysis. Nucleic Acids Res 36:W358–363. 10.1093/nar/gkn276

24. Wang M, Herrmann CJ, Simonovic M, et al (2015) Version 4.0 of PaxDb: Protein abundance data, integrated across model organisms, tissues, and cell-lines. Proteomics 15:3163–3168. 10.1002/pmic.201400441

25. Voskuhl RR, Itoh N, Tassoni A, et al (2019) Gene expression in oligodendrocytes during remyelination reveals cholesterol homeostasis as a therapeutic target in multiple sclerosis. Proc Natl Acad Sci U S A 116:10130–10139. 10.1073/pnas.1821306116

26. Liu Y, Tian F, Li S, et al (2021) Global effects of RAB3GAP1 dysexpression on the proteome of mouse cortical neurons. Amino Acids 53:1339–1350. 10.1007/s00726-021-03058-9

27. Oshlack A, Wakefield MJ (2009) Transcript length bias in RNA-seq data confounds systems biology. Biol Direct 4:14. 10.1186/1745-6150-4-14

28. Young MD, Wakefield MJ, Smyth GK, Oshlack A (2010) Gene ontology analysis for RNA-seq: accounting for selection bias. Genome Biol 11:R14. 10.1186/gb-2010-11-2-r14

29. Love MI, Huber W, Anders S (2014) Moderated estimation of fold change and dispersion for RNA-seq data with DESeq2. Genome Biol 15:550. 10.1186/s13059-014-0550-8

30. Padonou F, Gonzalez V, Provin N, et al (2022) Aire-dependent transcripts escape Raver2-induced splice-event inclusion in the thymic epithelium. EMBO Rep 23:e53576. 10.15252/embr.202153576

31. Langfelder P, Horvath S (2008) WGCNA: an R package for weighted correlation network analysis. BMC Bioinformatics 9:559. 10.1186/1471-2105-9-559

32. Mubeen S, Hoyt CT, Gemünd A, et al (2019) The Impact of Pathway Database Choice on Statistical Enrichment Analysis and Predictive Modeling. Front Genet 10:1203. 10.3389/fgene.2019.01203

33. Chen EY, Tan CM, Kou Y, et al (2013) Enrichr: interactive and collaborative HTML5 gene list enrichment analysis tool. BMC Bioinformatics 14:128. 10.1186/1471-2105-14-128

34. Simillion C, Liechti R, Lischer HEL, et al (2017) Avoiding the pitfalls of gene set enrichment analysis with SetRank. BMC Bioinformatics 18:151. 10.1186/s12859-017-1571-6

35. Maleki F, Ovens K, Hogan DJ, Kusalik AJ (2020) Gene Set Analysis: Challenges, Opportunities, and Future Research. Front Genet 11:654. 10.3389/fgene.2020.00654

36. Wu T, Hu E, Xu S, et al (2021) clusterProfiler 4.0: A universal enrichment tool for interpreting omics data. Innov Camb Mass 2:100141. 10.1016/j.xinn.2021.100141

37. Salka K, Abutaleb K, Chorvinsky E, et al (2021) IFN Stimulates ACE2 Expression in Pediatric Airway Epithelial Cells. Am J Respir Cell Mol Biol 64:515–518. 10.1165/rcmb.2020-0352LE

38. Wandel MP, Pathe C, Werner EI, et al (2017) GBPs Inhibit Motility of Shigella flexneri but Are Targeted for Degradation by the Bacterial Ubiquitin Ligase IpaH9.8. Cell Host Microbe 22:507-518.e5. 10.1016/j.chom.2017.09.007

39. Godugu K, Hay BA, Glinsky GV, Mousa SA (2023) Discovery of novel thyrointegrin αvβ3 antagonist fb-PMT (NP751) in the management of human glioblastoma multiforme. Neuro-Oncol Adv 5:vdac180. 10.1093/noajnl/vdac180

40. Crow M, Lim N, Ballouz S, et al (2019) Predictability of human differential gene expression. Proc Natl Acad Sci U S A 116:6491–6500. 10.1073/pnas.1802973116

41. Khatri P, Sirota M, Butte AJ (2012) Ten years of pathway analysis: current approaches and outstanding challenges. PLoS Comput Biol 8:e1002375. 10.1371/journal.pcbi.1002375

42. Leek JT, Storey JD (2007) Capturing heterogeneity in gene expression studies by surrogate variable analysis. PLoS Genet 3:1724–1735. 10.1371/journal.pgen.0030161

43. Zhang Y, Parmigiani G, Johnson WE (2020) ComBat-seq: batch effect adjustment for RNA-seq count data. NAR Genomics Bioinforma 2:qaa078. 10.1093/nargab/lqaa078

44. Fei T, Yu T (2020) scBatch: batch-effect correction of RNA-seq data through sample distance matrix adjustment. Bioinforma Oxf Engl 36:3115–3123. 10.1093/bioinformatics/btaa097

45. Lim N, Tesar S, Belmadani M, et al (2021) Curation of over 101000 transcriptomic studies to enable data reuse. Database J Biol Databases Curation 2021:baab006. 10.1093/database/baab006

46. Ashburner M, Ball CA, Blake JA, et al (2000) Gene ontology: tool for the unification of biology. The Gene Ontology Consortium. Nat Genet 25:25–29. 10.1038/75556

47. Kanehisa M, Goto S (2000) KEGG: kyoto encyclopedia of genes and genomes. Nucleic Acids Res 28:27–30. 10.1093/nar/28.1.27

48. Conesa A, Madrigal P, Tarazona S, et al (2016) A survey of best practices for RNA-seq data analysis. Genome Biol 17:13. 10.1186/s13059-016-0881-8

49. Dimitrieva S, Bucher P (2013) UCNEbase--a database of ultraconserved non-coding elements and genomic regulatory blocks. Nucleic Acids Res 41:D101–109. 10.1093/nar/gks1092

50. Zou Z, Ohta T, Miura F, Oki S (2022) ChIP-Atlas 2021 update: a data-mining suite for exploring epigenomic landscapes by fully integrating ChIP-seq, ATAC-seq and Bisulfite-seq data. Nucleic Acids Res 50:W175–W182. 10.1093/nar/gkac199

51. Mahi NA, Najafabadi MF, Pilarczyk M, et al (2019) GREIN: An Interactive Web Platform for Re-analyzing GEO RNA-seq Data. Sci Rep 9:7580. 10.1038/s41598-019-43935-8

52. Tarca AL, Bhatti G, Romero R (2013) A comparison of gene set analysis methods in terms of sensitivity, prioritization and specificity. PloS One 8:e79217. 10.1371/journal.pone.0079217

53. Guala D, Sonnhammer ELL (2017) A large-scale benchmark of gene prioritization methods. Sci Rep 7:46598. 10.1038/srep46598

54. Geistlinger L, Csaba G, Santarelli M, et al (2021) Toward a gold standard for benchmarking gene set enrichment analysis. Brief Bioinform 22:545–556. 10.1093/bib/bbz158

55. de Hoon MJL, Imoto S, Nolan J, Miyano S (2004) Open source clustering software. Bioinforma Oxf Engl 20:1453–1454. 10.1093/bioinformatics/bth078

56. Hunter JD (2007) Matplotlib: A 2D Graphics Environment. Comput Sci Eng 9:90–95. 10.1109/MCSE.2007.55

57. Waskom M (2021) seaborn: statistical data visualization. J Open Source Softw 6:3021. 10.21105/joss.03021

58. Wickham H (2016) ggplot2: elegant graphics for data analysis, Second edition. Springer international publishing, Cham

